# Kynurenine metabolism is altered in *mdx* mice: a potential muscle to brain connection

**DOI:** 10.1101/2022.01.14.476359

**Authors:** Emily N. Copeland, Kennedy C. Whitley, Colton J.F. Watson, Bradley J. Baranowski, Nigel Kurgan, Adam J. MacNeil, Rebecca E.K. MacPherson, Val A. Fajardo, David J. Allison

## Abstract

Regular exercise can direct muscle kynurenine (KYN) metabolism toward the neuroprotective branch of the kynurenine pathway thereby limiting the accumulation of neurotoxic metabolites in the brain and contributing to mental resilience. While the effect of regular exercise has been studied, the effect of muscle disease on KYN metabolism has not yet been investigated. Previous work has highlighted anxiety-like behaviors in approximately 25% of patients with DMD, possibly due to altered KYN metabolism. Here, we characterized KYN metabolism in *mdx* mouse models of Duchenne muscular dystrophy (DMD). Young (8-10 week old) DBA/2J (D2) *mdx* mice, but not age-matched C57BL/10 (C57) *mdx* mice, had lower levels of circulating KYNA and KYNA:KYN ratio compared with their respective wild-type (WT) controls. Moreover, only D2 *mdx* mice displayed signs of anxiety-like behaviour, spending more time in the corners of their cages during a novel object recognition test when compared with WT. Along with this, we found that muscles from D2 *mdx* mice had less peroxisome proliferator-activated receptor-gamma coactivator 1-alpha and kynurenine amino transferase-1 enzyme content as well as elevated expression of inflammatory cytokines compared with WT muscles. Thus, our pilot work shows that KYN metabolism is altered in D2 *mdx* mice, with a potential contribution from altered muscle health.

## Introduction

Kynurenine (KYN) is a metabolite of the amino acid L-tryptophan and alterations in its plasma levels have been strongly correlated with depression and anxiety (Ogyu et al. 2018). KYN can be metabolized peripherally, or within the brain, following passage across the blood brain barrier, along one of two distinct branches. The KYN-KYNA branch is regulated by the enzyme kynurenine aminotransferase (KAT) and is considered neuroprotective as it degrades KYN into the non-blood brain barrier (BBB) transportable metabolite kynurenic acid (KYNA). The KYN-NAD branch is regulated by the enzyme kynurenine monooxygenase (KMO) and is considered neurotoxic as it degrades KYN into the BBB transportable metabolite 3-hydroxykynurenine (3-HK) and, further in the cascade, quinolinic acid (QUIN), both of which can stimulate free radical production and neuronal apoptosis (Allison and Ditor 2014, 2015). Interestingly, there is a muscle-to-brain connection, with muscle-specific expression of KAT directing KYN metabolism towards the neuroprotective KYN-KYNA branch (Allison et al. 2019). In rodents and humans, regular exercise can increase the expression of KAT via increased peroxisome proliferator-activated receptor-gamma coactivator 1-alpha (PGC-1α) expression/activation and a reduction in inflammation, ultimately leading to increased production of KYNA from KYN (Agudelo et al. 2019; Agudelo et al. 2014). While research has shown that promoting muscle health through exercise can influence mental health through this pathway (Agudelo et al. 2014), it is unclear as to whether diseases that specifically affect muscle can negatively impact this pathway, ultimately contributing to anxiety and depression.

Duchenne muscular dystrophy (DMD) is a severe X-linked progressive muscle wasting disease caused by mutations in the dystrophin gene that lead to the complete absence of dystrophin protein. Normally required to provide membrane stability in cardiac and skeletal muscle, the absence of dystrophin results in muscle fragility, leading to extensive degeneration/regeneration cycling, weakness, and fatigue (Bulfield et al. 1984; Dangain and Vrbova 1984). With no cure for DMD, patients will have a shortened lifespan due to cardiorespiratory complications. In addition to muscular deficits, approximately one third of patients also experience cognitive difficulties, such as lowered IQs, impaired memory, and reading difficulties (Billard et al. 1992; Cotton et al. 2001). Studies have also shown that psychiatric disturbances, such as depression and anxiety, are common among patients living with DMD (Fitzpatrick et al. 1986; Roccella et al. 2003) - further reducing quality of life. Although the many physical and psychosocial hardships experienced by those living with DMD are major contributors to depression and anxiety, unfavourable alterations in the kynurenine pathway may also contribute to these symptoms. The extent to which KYN metabolism is altered in muscular dystrophy, and its relationship to depression and anxiety remains unknown.

In the present study, we characterized anxiety-like symptoms and KYN metabolism in *mdx* mouse models of DMD. We utilized young 8-10 week old DBA/2J (D2) *mdx* mice and C57BL/10 (C57) *mdx* mice as they are known to differ in disease severity, with the D2 *mdx* mice presenting with earlier-onset and worsened pathology (Coley et al. 2016; Hammers et al. 2020; van Putten et al. 2019). As such, we hypothesized that KYN metabolism would be negatively impacted in young D2 *mdx* mice further producing anxiety-like behaviours, more so than age-matched C57 *mdx* mice.

## Methods

### Animals

Male C57 *mdx*, C57 WT, D2 *mdx*, and D2 WT mice (n = 6 group) were purchased at 6-7 weeks of age from Jackson Laboratories. The mice were acclimated and housed in Brock University’s Animal Facility, in an environmentally controlled room with a standard 12:12 hour light-dark cycle. The mice were allowed access to food and water *ad libitum*. After the mice reached 8-10 weeks of age, they were euthanized via exsanguination under general anesthetic (vaporized isoflurane) and their muscles and serum were collected. All animal procedures were reviewed and approved by the Animal Care Committee of Brock University and followed the guidelines of the Canadian Council on Animal Care (AUP 17-06-03).

### Serum collection and tissue dissection

Extensor digitorum longus (EDL), red gastrocnemius (RG), white gastrocnemius (WG), diaphragm, and serum samples were collected from euthanized animals and flash frozen in liquid nitrogen. For serum, blood was collected and then spun for 5 min at 8,000 × g prior to collecting the supernatant. The serum and muscle samples were kept frozen at −80°C until experimentation.

### ELISA

Serum KYN and KYNA was measured using the commercially available L-Kynurenine ELISA (ImmuSol, BA-E-2200) and KYNA ELISA (ImmuSol, IS-I-0200) kits, respectively. All samples and controls were measured in duplicate. Absorbance of each plate was measured at 450 nm, with a reference wavelength of 540 nm using an M2 Molecular Devices MultiMode plate reader.

### Western blotting

Western blotting was performed on the EDL of D2 WT and *mdx* mice to determine protein concentration of PGC-1α, KAT1, and KAT3. Protein from muscle homogenates were solubilized in 4X Laemmli buffer (BioRad, 161-0747 final concentration 1x) and was and then electrophoretically separated at 240V for 22 minutes on a 7-12% TGX gradient gel (BioRad, 4568081). Proteins were then transferred to a PVDF membrane using a BioRad Trans Blot Turbo. Subsequently, membranes were blocked for 1 hour in a 5% (w/v) milk and Tris-buffered saline tween (TBST) solution. Primary antibodies (1:2000 dilution) were then added and membranes were incubated overnight at 4°C. Following incubation, membranes were washed three times using TBST, then incubated with corresponding secondary antibodies (goat anti-mouse, Novus, MAB7947; donkey anti-rabbit, PAS, 68622; all at 1:2000) at room temperature for one hour. Membranes were then washed again before chemiluminescent substrate Milipore Immobilon (WBKLS0500; Sigma-Aldrich) was added prior to imaging with a BioRad Chemidoc. Optical densities were analyzed with ImageLab (BioRad) and normalized to GAPDH (ProteinTech, 60004; 1:5000).

### mRNA analysis

WG, RG, and diaphragm were homogenized with 1 mL of TRIzol. A Qiagen RNeasy kit spin column was used to isolate RNA from the muscle samples, then quantified using a Nanoview Plus spectrophotometer. EcoDry RNA to cDNA reaction tubes and a SimpliAmp Thermal Cylinder were then used to facilitate the generation of cDNA which was then analyzed using a 96-well qPCR reaction plate and the StepOnePlus Real-Time PCR System (Applied Biosystems, #272004476). Assays were performed with KAPA SYBR FAST (Roche, Diagnostics) mastermix and amplification efficiency-optimized primers: IL-6 (AGACAAAGCCAGAGTCCTTCAGAGA forward and TGGTCTTGGTCCTTAGCCACTCC reverse); TNF-(TGAACTTCGGGGTGATCGGTCC forward and TCCAGCTGCTCCTCCACTTGGT reverse); and GAPDH (CGGTGCTGAGTATGTCGTGGAGTC forward and GGGGCTAAGCAGTTGGTGGTG reverse) (IDT). Threshold cycle (Ct) values were recorded and data was analyzed using the ΔΔCt method, using expression of GAPDH as a reference gene.

### Analysis of Anxiety-Like Behaviour

A novel object recognition test (NORT) was performed in an arena containing four equal sized, open top boxes, over three 10 minute phases: acclimatization, training, and testing (Hayward et al. 2022). Pilot testing during the acclimation period indicated no preference to any specific locations within each box. Movement was recorded with a video camera, secured above the arena across 10 min testing sessions. Videos were examined manually by a blind experimenter for exploration of novel and familiar objects, total movement time, and time spent in corners. The exploration times were reduced only in the D2 *mdx* mouse and was not due to differences in time spent moving between the C57 and D2 *mdx* mice (Hayward et al. 2022). However, further analysis of specific locations during mice movement was not analyzed. In an open field test, anxiety-like behaviour is described as time spent in the corners or periphery of the arena (La-Vu, 2020). Therefore, as an anxiety-like measure, we compared the total elapsed time spent in the corners of the box to total elapsed time exploring the other areas of the box over a ten-minute period.

### Statistics

All data are expressed as mean ± SEM. Student’s *t* test was used for all comparisons using GraphPad Software with statistical significance and statistical trends set to *p* < 0.05 and *p* < 0.15, respectively.

## Results and Discussion

First, we examined KYN, KYNA and the KYNA:KYN ratio in serum samples obtained from the *mdx* and WT mice. Figure 1A shows that although serum concentration of KYN was not significantly different between D2 *mdx* and WT mice, the serum concentration of KYNA was significantly decreased in D2 *mdx* compared to D2 WT mice. This therefore produced a significant decrease in the ratio of KYNA:KYN in the serum of the D2 *mdx* mice compared to the D2 WT (Figure 1A). Although there was a significant reduction in the serum concentration of KYN in C57 *mdx* mice compared to C57 WT mice, significant decreases were not seen in serum concentration of KYNA, and no significant differences were found in the ratio of KYNA:KYN between groups (Figure 1B). Altogether, these results suggest that KYN metabolism is altered in D2 *mdx* mice and is perhaps reflective of a diversion away from the neuroprotective pathway.

**Figure 1.**
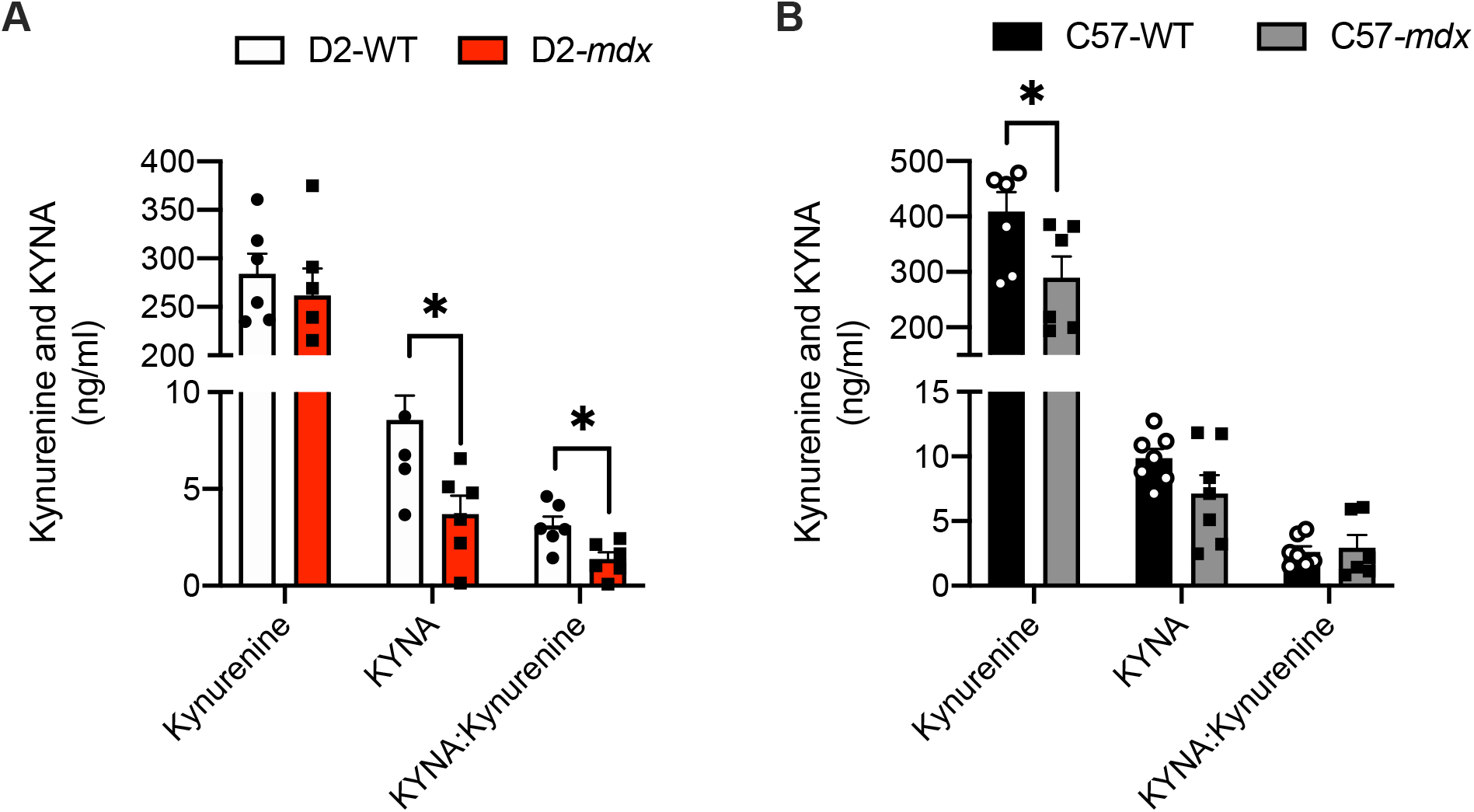
ELISA results of serum KYN and KYNA concentrations. A) Concentration of KYN did not significantly differ between D2 *mdx* and WT mice. D2 *mdx* mice had significantly reduced concentrations of KYNA (p < 0.05) and a significantly smaller KYNA:KYN ratio (p < 0.05) compared to D2 WT mice. B) C57 *mdx* mice had significantly less KYN concentration (p < 0.05) than C57 WT mice but, no significant differences were observed between C57 *mdx* and WT mice with regards to KYNA concentration or KYNA:KYN ratio. \**p*<0.05 using a Student’s t-test (n < 4-6 per group).

**Figure 2.**
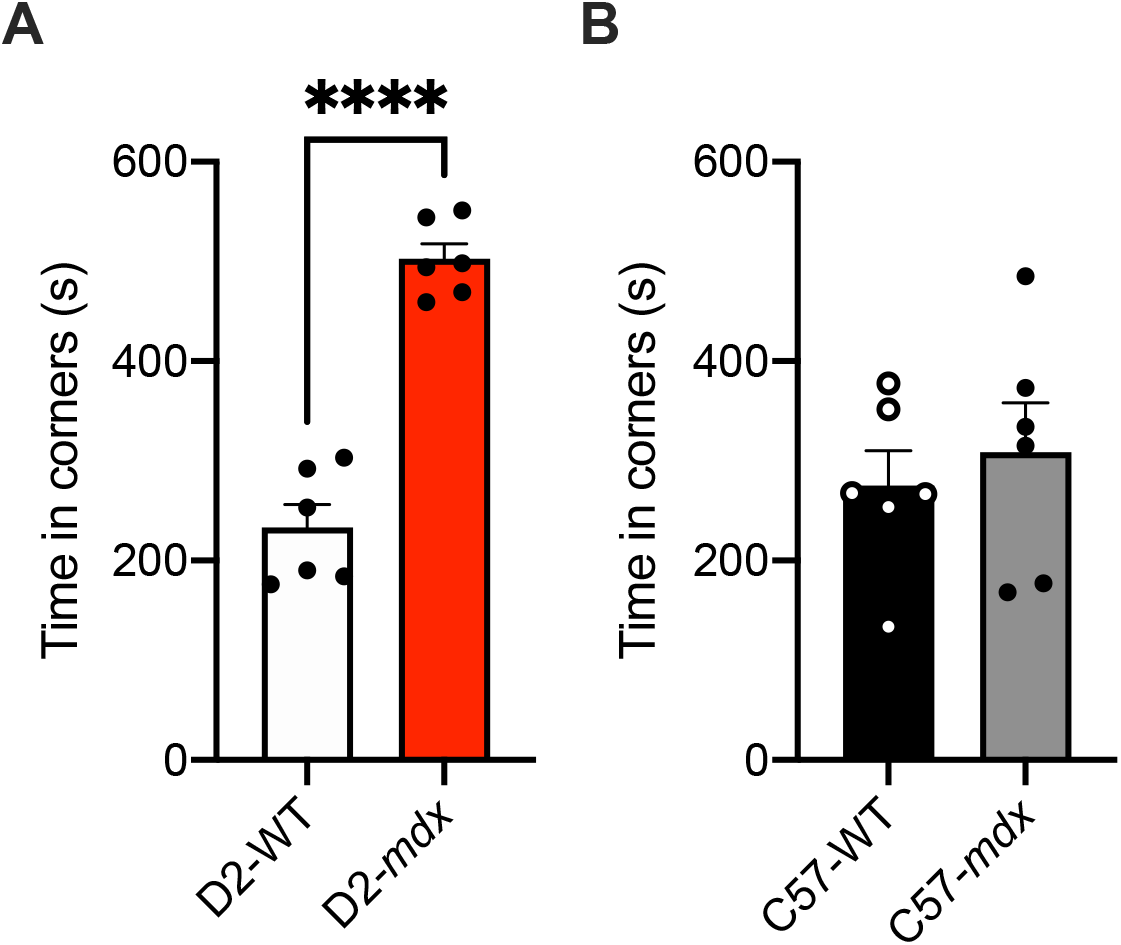
Time spent in corners of arena during novel object recognition test. A) D2 *mdx* mice spent significantly more time (p < 0.05) in the corners of the arenas than the D2 WT mice. B) C57 *mdx* and WT mice spent similar amounts of time in corners of the arenas. \*\*\*\**p*<0.0001 using a Student’s t-test (n = 4-6 per group).

The NORT is a commonly used behavioural test for rodents, specifically during memory-related tasks. It is expected that memory-intact rodents will explore a novel object for longer durations than a familiar object, while memory-impaired rodents will have no preference for either object, exploring both for similar durations (Ennaceur and Delacour 1988). In a recent study with the same mice used in this study, it was found that D2 *mdx* mice had significant memory deficits compared to their WT counterparts with significantly lower exploration time (Hayward et al. 2021). However, this was not observed in age-matched C57 *mdx* mice and we believe that this corresponds to the known differences in disease severity between the strains of mice (Coley et al. 2016; Hammers et al. 2020; van Putten et al. 2019). Additionally, the NORT can highlight anxiety-like behaviours in which rodents spend more time in the corners rather than in the center of the arena or exploring (Ennaceur et al. 2005). Our NORT showed that only D2 *mdx* mice spent more time in corners compared to the D2 WT mice; however, this difference was not found between C57 *mdx* and WT mice (Figure 4). This suggests that at 8-10 weeks of age, D2 *mdx* mice present with anxiety-like behaviour whereas *C57 mdx* mice may not. Our results are not consistent with those reported recently by Bagdatlioglu and colleagues (2020) who found that C57 *mdx* mice indeed spent more time in the corners compared with WT mice during a NORT (Bagdatlioglu et al. 2020). Explaining this discrepancy is the factor of age, since Bagdatlioglu et al., utilized 4, 12 and 18 month old mice; and in addition to finding an effect of *mdx*, they also reported an effect of aging, where the time spent in corners increased with increasing age in *mdx* mice. Thus, it is possible that the reason why we did not find any anxiety-like behaviours in young C57 *mdx* is because the disease had not progressed far enough. In contrast, we found anxiety-like behaviour in age-matched D2 *mdx* mice who are known to present with worsened and earlier onset dystrophic disease; and this result appears to coincide with altered KYN metabolism.

**Figure 3.**
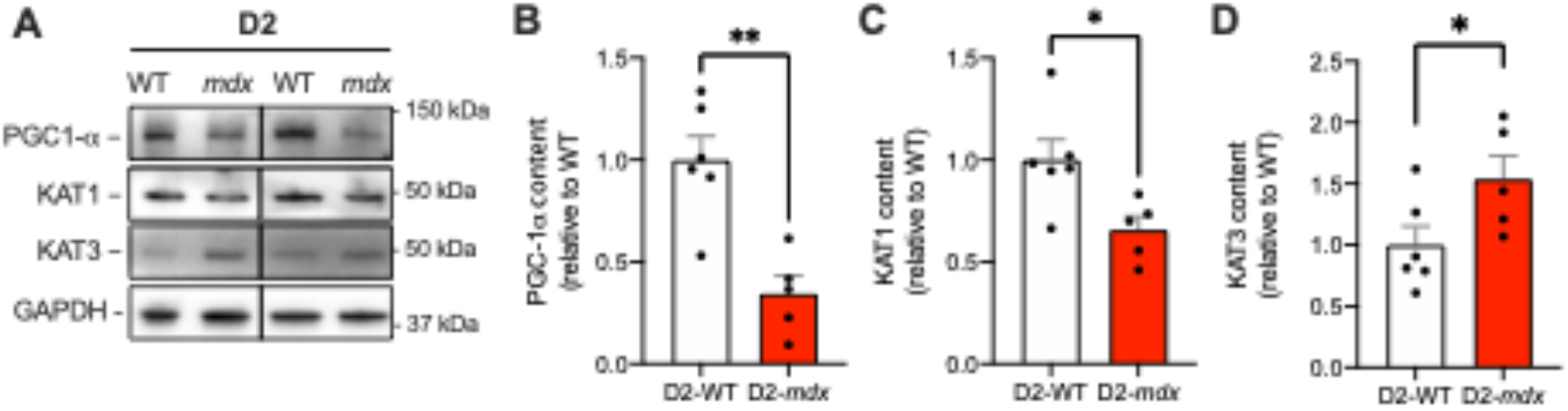
Western blot analysis of PGC-1α, KAT1, and KAT3 in EDL of D2 *mdx* and WT mice. A) Representative Western blot images. B) PGC-1α content was significantly reduced in D2 *mdx* mice compared to D2 WT mice. C) KAT1 content was significantly reduced in D2 *mdx* mice compared to WT mice. D) KAT3 content was significantly increased in D2 *mdx* mice compared to WT mice. \*\**p*<0.01, \**p*<0.05, using a Student’s t-test (n = 4-6 per group). All data were normalized to GAPDH.

**Figure 4.**
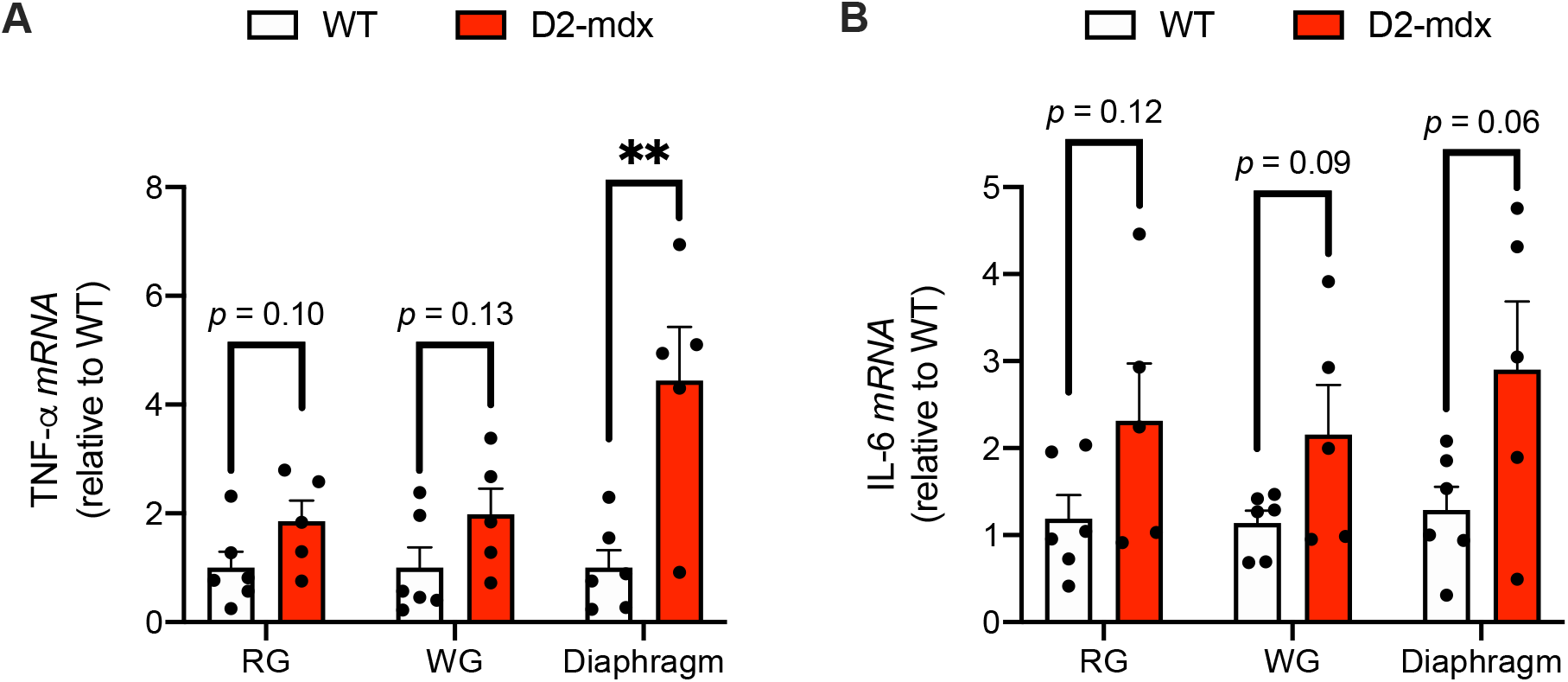
mRNA analyses for inflammatory markers TNF-α and IL-6 in D2 *mdx* and WT muscles. A) TNF-α mRNA content in red gastrocnemius (RG), white gastrocnemius (WG) and diaphragm muscles. B) IL-6 mRNA content increase in RG, WG, and diaphragm muscles from D2 *mdx* mice compared to the WT mice. \*\**p*<0.01, using a Student’s t-test (n = 5-6 per group).

To investigate the potential mechanisms leading to these alterations in KYN metabolism in D2 *mdx* mice we examined muscle PGC-1α and KAT expression. Our results show that PGC-1α is lowered in EDL muscles from D2 *mdx* mice compared with D2 WT mice (Figure 3A and B). Similarly, we saw a significant reduction in KAT1 content in D2 *mdx* mice compared to D2 WT mice (Figure 3B) but, the opposite was found of KAT3 content (Figure 3C). Interestingly, others have shown that in *Kat2*^−/−^ knockout mice, an isoform not found in skeletal muscle (Agudelo, Femenia, et al., 2014), both KAT1 and KAT3 were upregulated in the brains of these mice as a way to compensate for the lack of KAT2 (Yu et al. 2006). Similarly, we speculate that the increase in KAT3 may be a failed compensatory mechanism (for the loss of KAT1) aimed at increasing the KYNA:KYN ratio; and the mechanisms explaining this upregulation of KAT3 are independent of PGC-1α and should be explored with future research. In addition, Yu et al., (2006) found that KAT3 was highly expressed in murine cardiac muscle, which is solely composed of type I muscle fibres. Since we have found that EDL muscles from D2 *mdx* mice display an increased presence of slow fibers (data not shown), it is possible that the increase in KAT3 may also be reflective of a fibre type shift.

In addition to exploring PGC-1α and KAT content, we also assessed levels of muscle inflammation. Across RG, WG and diaphragm samples, we found elevated levels of TNF-α and IL-6 in D2 *mdx* mice compared to WT (Figure 4), however, most comparisons were only trending towards significance – likely limited in sample size and variability. Despite the general lack of statistical significance, these findings are congruent with those from a previous study showing elevated pro-inflammatory cytokine expression in young (8-10 week old) D2 *mdx* mice (Coley et al. 2016); and we speculate that heightened inflammation in these mice can also contribute to the alterations in KYN metabolism as they preferentially upregulate indoleamine 2,3 dioxygenase (IDO), the initiator of KYN metabolism, and KMO (Campbell et al. 2014), ultimately favouring the neurotoxic branch.

To our knowledge, ours is the first study to show that KYN metabolism is altered in the D2 *mdx* mouse, a result that is associated with anxiety-like behaviour. The D2 *mdx* mouse has only recently emerged as a viable model for DMD, and future studies should examine whether targeting KYN metabolism by improving muscle health can attenuate anxiety in these mice, ultimately improving their quality of life. Physical activity is well known to influence KYN metabolism, and low-intensity treadmill exercise has been shown to promote an oxidative phenotype shift and PGC-1α expression in *mdx* mice (Frinchi et al. 2021), making it a viable therapeutic option that could be investigated further. In humans with spinal cord injuries, consuming an anti-inflammatory diet also positively alters KYN metabolism and markers of mental health (Allison and Ditor 2018); and it would thus be of interest to determine whether an anti-inflammatory diet could also benefit those living with DMD.

While this initial preliminary work will likely lead to future studies aimed at targeting kynurenine metabolism in *mdx* mice and possibly patients with DMD, we do acknowledge a few limitations. First, small sample volumes prevented our analyses of neurotoxic metabolites such as QUIN. Secondly, future research of KYN metabolism in *mdx* mice may benefit from larger sample sizes across young and old-aged groups to equally distribute sample across all KYN metabolism branches. Finally, it would also be of interest to determine whether KYN metabolism is altered in patients with DMD, as this would ultimately set the stage for future studies aimed at altering this pathway in these patients.

In conclusion, our study provides novel insight onto the anxiety and depressive-like symptoms found in *mdx* mice, perhaps suggestive of a muscle-to-brain connection. This study provides rationale for future studies examining whether targeting KYN metabolism in DMD and *mdx* muscle through genetic or pharmacological/nutraceutical approaches can improve upon these outcomes in the preclinical and clinical setting.

## Acknowledgements

This work was funded by a Canada Research Chair Tier 2 Award to VAF.

